# Nociception in chicken embryos, Part I: Analysis of cardiovascular responses to a mechanical noxious stimulus

**DOI:** 10.1101/2023.04.14.536899

**Authors:** Larissa Weiss, Anna M. Saller, Julia Werner, Stephanie C. Süß, Judith Reiser, Sandra Kollmansperger, Malte Anders, Thomas Fenzl, Benjamin Schusser, Christine Baumgartner

## Abstract

While it is assumed that chicken embryos acquire the ability for nociception during the developmental period in the egg, an exact time point has not yet been specified. This study aimed to determine the onset of nociception during embryonic development in chicken. Changes in blood pressure and heart rate (HR) in response to a mechanical noxious stimulus at the base of the beak versus a light touch on the beak in chicken embryos between embryonic days (EDs) 7 and 18 were examined. Mean arterial pressure (MAP) was the most sensitive parameter for assessing cardiovascular responses. Significant changes in MAP in response to a noxious stimulus were detected in embryos at ED16 to ED18, while significant changes in HR were observed on ED17 and ED18. Infiltration anesthesia with the local anesthetic lidocaine significantly reduced reactions in MAP on ED18, so the cardiovascular changes can be assumed to be nociceptive responses.

## Introduction

In present times animal welfare has increasingly become the focus of public attention with regard to farm and laboratory animals. Consequently, the culling of male day-old chicken for economic reasons is more and more ethically questioned. A large part of the male offspring in the layer industry is killed after hatching, as the fattening of male layer type chicken is not economically profitable^1^. In the EU, 330 million male chicks are killed annually through maceration or gasification^2^, which is currently evoking a major discussion. Germany and France have already adapted their laws and banned the killing of male day-old chicks for economic reasons although there is not yet EU-wide regulation^3^. As an alternative, *in ovo* sex determination with subsequent killing of male embryos is already being practiced^4^. However, it is important for animal welfare reasons and for the public acceptance of *in ovo* sex determination that related culling be conducted at an early stage of development when nociception and the perception of pain are not yet possible^4,5^.

Furthermore, chicken embryos are of great importance for biomedical research because of advantages provided in terms of fast growth and good accessibility in various research areas, such as developmental biology, toxicology, cancer research and drug development^6,7^. Under European regulations, interventions and treatments on chicken embryos are not considered animal experiments and even count as a replacement method in the context of the 3R principles^8^. At this time, there are no regulations regarding anesthesia and analgesia of chicken embryos during painful interventions^6,8^. Greater clarity regarding the period during which chicken embryos are capable of nociception and pain sensation would lead to improved animal welfare in research.

In pain research, a fundamental distinction is made between nociception and the perception of pain^9^. While nociception is the detection of a potentially tissue-damaging stimulus and its transmission by the nociceptive nervous system^10,11^, pain is characterized by a subjective, conscious sensory perception, usually triggered by nociception^12,13^. Nociception and pain are progressive adaptive processes that gradually develop throughout the fetal period^14^. It is considered confirmed, that the chicken embryo acquires the ability for nociception at some point during the 21-day developmental period in the egg^8,15^. However, the question at which exact time point nociception or even pain sensation can be presumed is controversial. In several publications, researchers agree that nociception and pain perception are not possible in the first trimester of embryonic development of the chicken^4,15^. A requirement for the ability to perceive pain is the existence of functional pathways that enable the transmission of stimuli to the brain^12,14^. Although the first sensory afferent nerve fibers develop on embryonic day (ED) 4, the closure of multisynaptic reflex arcs does not occur until ED7^16-18^. It is described in the literature that the chicken embryo develops a functional brain on ED13^15,19^. However, it is only confirmed that the brain does not show any electrical activity until 6.5 days of incubation^20^. Pain sensation is therefore considered to be impossible up to ED7, but beyond that, no specific time point can be defined from which the chicken embryo is capable of nociception and pain sensation^4,15^.

Since self-reporting, which is the gold standard in humans to detect pain^21^, is not possible as a direct method of pain evaluation in animals, indirect methods such as alteration of physiological and behavioral parameters must be resorted to^22^. Changes in heart rate and blood pressure are therefore used as clinical indicators of nociception and pain^23^.

This study is part of a comprehensive study in which the nociceptive ability of chicken embryos was investigated using cardiovascular parameters, behavioral observations and EEG. Here, we present the results of the cardiovascular study and, in particular, the implemented cardiovascular measurement methods in regard to chicken embryos that were designed for investigation of the time point at which chicken embryos are able to respond to a noxious stimulus with a nociceptive cardiovascular response. The corresponding results of the EEG measurements and behavioral observations and the implemented techniques will be presented in further publications.

## Results

### Increasing arterial pressure and evolution of heart rate during embryonic development of the chicken

Systolic (SAP), diastolic (DAP) and mean arterial pressure (MAP) in the chorioallantoic artery and heart rate (HR) were recorded over one minute at ED7, ED9 and EDs 12 to 18. SAP, DAP and MAP increased with age of the embryos (Table 1). ED7 showed the lowest MAP with a value of 2.08 mmHg ± 0.40, and ED18 showed the highest MAP with a value of 17.28 mmHg ± 3.04.

**Table 1.** SAP, DAP, MAP and HR at ED7 (n=3), ED9 (n=6), and ED12 to ED18 (n=10). Values are shown as the mean ± standard deviation.

|  | ED7 | ED9 | ED12 | ED13 | ED14 | ED15 | ED16 | ED17 | ED18 |
| --- | --- | --- | --- | --- | --- | --- | --- | --- | --- |
| SAP<br>(mmHg) | 3.50<br>$\pm 0.65$ | 6.04<br>$\pm 1.46$ | 9.19<br>$\pm 1.32$ | 9.88<br>$\pm 1.52$ | 13.02<br>$\pm 1.60$ | 16.54<br>$\pm 3.04$ | 21.44<br>$\pm 2.78$ | 24.46<br>$\pm 5.50$ | 24.65<br>$\pm 4.36$ |
| DAP<br>(mmHg) | 1.07<br>$\pm 0.36$ | 1.98<br>$\pm 1.10$ | 2.20<br>$\pm 1.12$ | 2.96<br>$\pm 0.61$ | 3.95<br>$\pm 1.14$ | 5.69<br>$\pm 1.82$ | 7.80<br>$\pm 2.16$ | 10.77<br>$\pm 3.53$ | 11.43<br>$\pm 2.43$ |
| MAP<br>(mmHg) | 2.08<br>$\pm 0.40$ | 3.44<br>$\pm 1.24$ | 4.83<br>$\pm 1.05$ | 5.52<br>$\pm 0.79$ | 7.32<br>$\pm 1.26$ | 10.11<br>$\pm 2.45$ | 13.73<br>$\pm 2.38$ | 16.79<br>$\pm 4.21$ | 17.28<br>$\pm 3.04$ |
| HR<br>(bpm) | 128.97<br>$\pm 15.40$ | 147.57<br>$\pm 9.03$ | 159.08<br>$\pm 26.64$ | 146.61<br>$\pm 19.99$ | 179.10<br>$\pm 35.06$ | 154.33<br>$\pm 33.12$ | 151.35<br>$\pm 36.44$ | 179.08<br>$\pm 29.18$ | 176.07<br>$\pm 35.75$ |

### Increase in MAP in response to a noxious stimulus

The response in MAP to a noxious mechanical stimulus at the base of the beak (*Pinch*) was compared to a light touch at the base of the beak as a negative control (*Touch*) in embryos between EDs 7 and 18. As shown in Fig. 1, a significant increase in MAP was observed as a reaction to *Pinch* in embryos at ED16 (p=0.0008), ED17 (p=0.0020) and ED18 (p=0.0048). ED18 embryos showed the strongest response in MAP, with an increase of 15.52 % ± 12.36 from baseline. In comparison, a deviation from baseline of only 1.30 % ± 0.94 was detected in response to *Touch* on ED18. In embryos at ED7, ED9 and EDs 12 to 15, no significant changes in MAP in response to *Pinch* compared to *Touch* were detected, which can be seen in Supplementary Fig. 1 and Supplementary Fig. 2.

**Fig 1.**
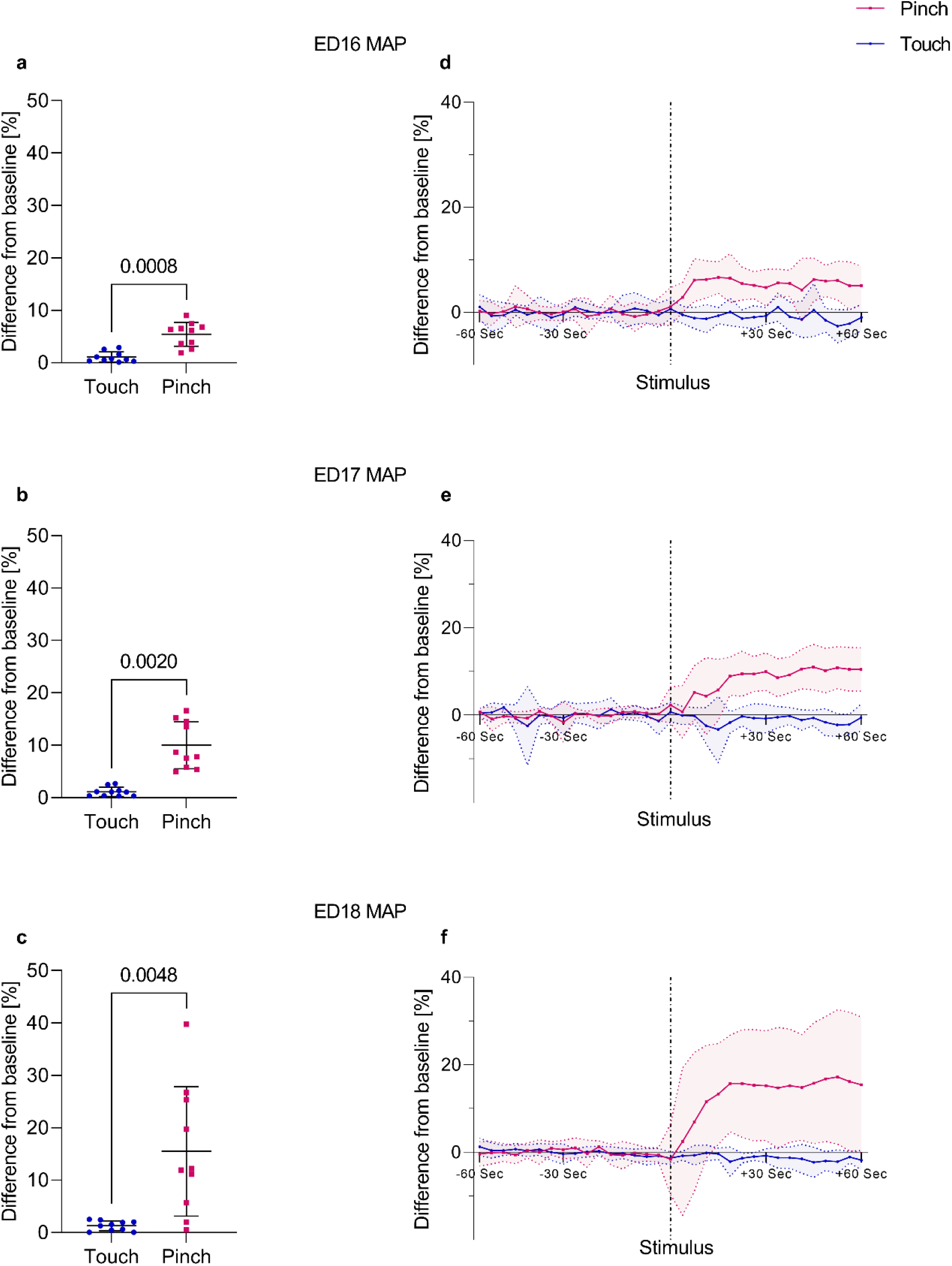
Percent change in MAP post *Touch* and *Pinch*. Embryos at EDs 16 to 18 (n=10) received a mechanical noxious stimulus (*Pinch*) and a light touch as a negative control (*Touch*) at the base of the beak in randomized order. **a-c** Percent change to baseline in MAP post *Pinch* compared to *Touch*. Displayed as the mean ± standard deviation. Paired t test (normally distributed: a and c) or Wilcoxon signed-rank test (not normally distributed: b). P values shown; a: p=0.0008, b: p=0.0020, c: p=0.0048. **d-f** Percent change from the baseline mean value in MAP over time; values recorded every four seconds for one minute before and one minute after stimulation (*Pinch* and *Touch*); values shown as the mean ± standard deviation (shaded).

### Changes in HR up to a noxious stimulus

Regarding HR, two reaction patterns have been observed, particularly in ED17 and ED18 embryos. In some embryos, HR immediately increased post *Pinch*. In other embryos a hyperacute decrease in HR followed by an increase was observed in response to *Pinch*, as shown in Fig. 2. A decrease in HR of at least -15 % with a subsequent increase of at least 5 % from the baseline mean value post *Pinch* was observed in 80 % of ED18 embryos and in 30 % of ED17 embryos and was not detected post *Touch*. In embryos at ED18 HR decreased by up to -48.54 % ± 19.71 over 9.50 s ± 6.02 on average post *Pinch*. At ED17 these embryos showed a decrease in HR by up to -41.87 % ± 8.32 over 16.00 s ± 6.93 on average post *Pinch*. Simultaneous to the hyperacute decrease in HR, a slight decrease in MAP was also observed, particularly when the decrease in HR was large. In embryos at ED15 and ED16, the observations were inconsistent and could not be clearly distinguished from physiological variations in HR. In younger embryos, no hyperacute decrease in HR was observed in response to *Pinch*.

**Fig 2.**
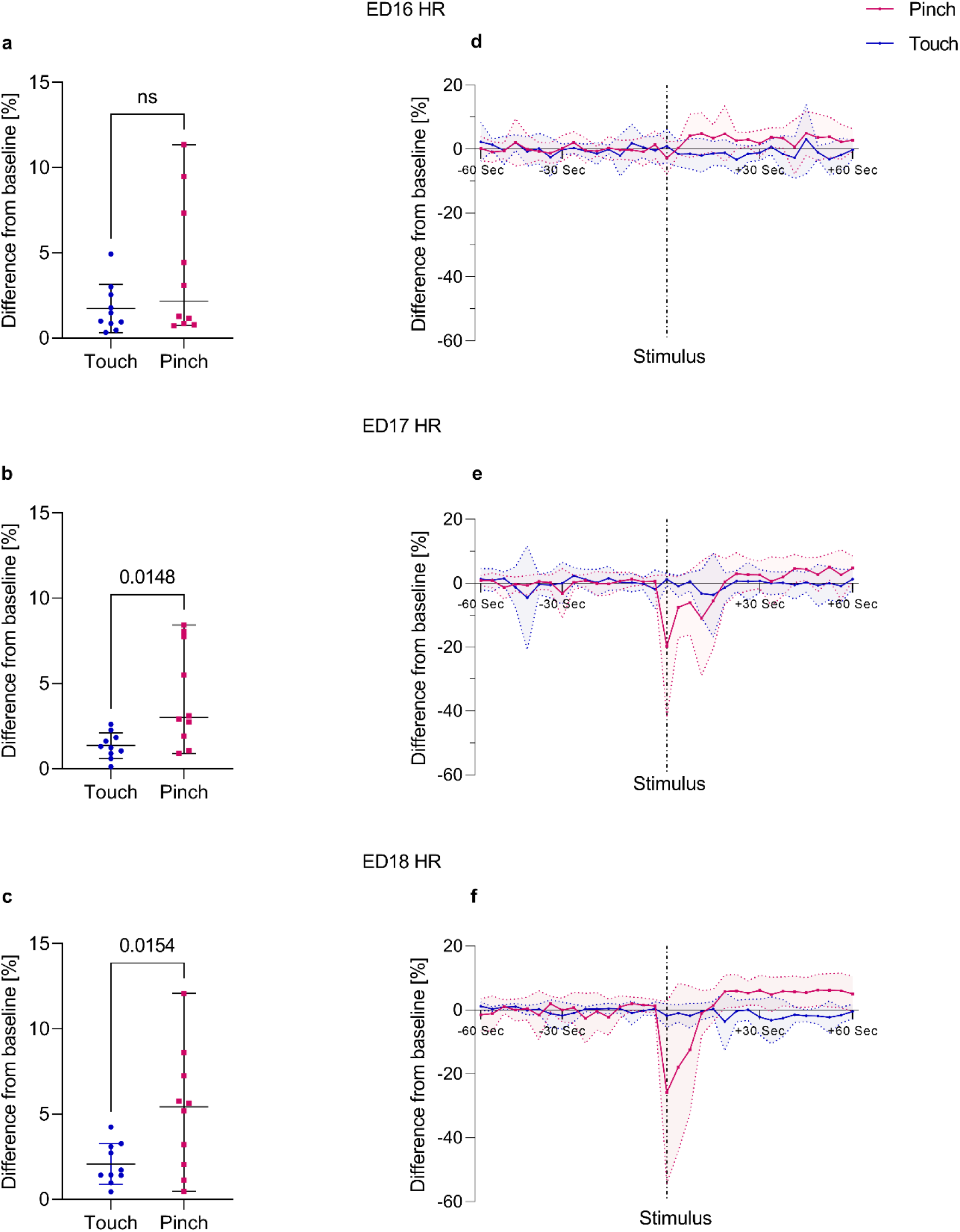
Percent change in HR post *Touch* and *Pinch*. Embryos at EDs 16 to 18 (n=10) received a mechanical noxious stimulus (*Pinch*) and a light touch as a negative control (*Touch*) at the base of the beak in randomized order. **a-c** Percent change to baseline in HR post *Pinch* compared to *Touch*. Displayed as the mean ± standard deviation. Paired t test (normally distributed: b and c) or Wilcoxon signed-rank test (not normally distributed: a). P values shown; b: p=0.0148, c: p=0.0154; ns=no significant difference between the groups (a). **d-f** Percent change from the baseline mean value in HR over time; values recorded every four seconds for one minute before and one minute after stimulation (*Pinch* and *Touch*); values shown as the mean ± standard deviation (shaded).

Significant increases in HR in response to *Pinch* compared to *Touch* were detected in embryos at ED17 (p=0.0148) and ED18 (p=0.0154) (Fig. 2). Embryos at ED18 showed the largest increase in HR post *Pinch* with a deviation of 5.14 % ± 3.60 from baseline, compared to a deviation of only 2.07 % ± 1.20 from baseline post *Touch*. At ED7, ED9 and EDs 12 to 16, no significant changes in HR were observed, which can be seen in Supplementary Fig. 3 and Supplementary Fig. 4.

### Reduction of cardiovascular response by local anesthesia

The application of the local anesthetic lidocaine (*Lido*) at the base of the beak prior to stimulation significantly reduced the MAP increase in response to *Pinch* in embryos at ED18. Compared to the group without local anesthesia (*ED18 w/o Lido*), which showed an increase of 15.52 % ± 12.36 post *Pinch*, the increase in MAP was reduced to 5.00 % ± 3.42 in the group that received lidocaine (*ED18 w/ Lido*). As represented in Fig. 3, the *ED18 w/o Lido Pinch* group showed the largest increase in MAP in response to *Pinch* compared to *ED18 w/o Lido Touch* (p=0.0007), *ED18 w/ Lido Touch* (p=0.0031) and *ED18 w/ Lido Pinch* (p=0.0397).

**Fig 3.**
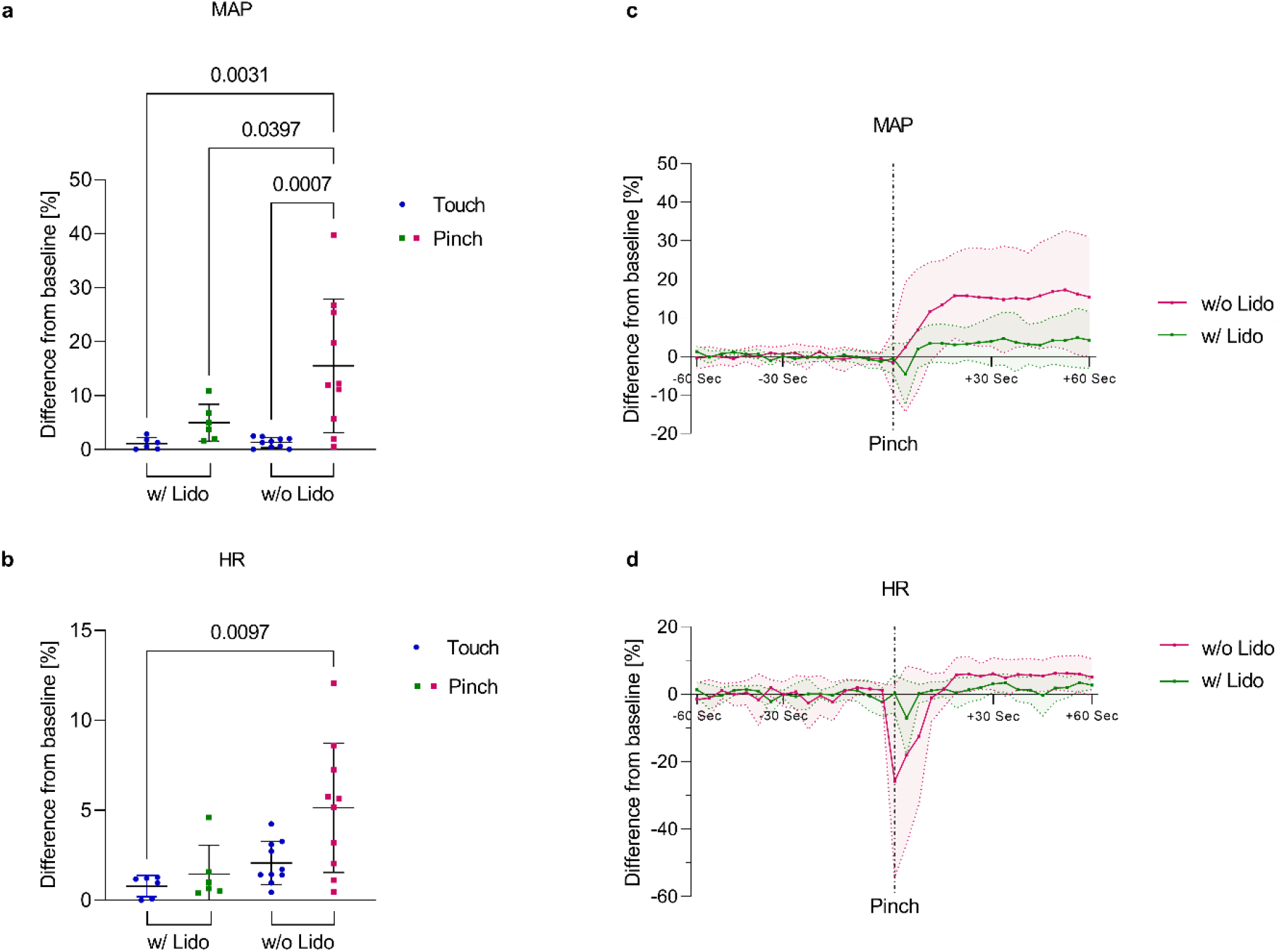
Local anesthesia control group. Percent change in MAP and HR post *Touch* and *Pinch*. ED18 embryos either received a lidocaine injection (*ED18 w/ Lido*; n=6) or no lidocaine injection (*ED18 w/o Lido*; n=10) at the base of the beak prior to stimulation (*Touch* and *Pinch*). **a** Percent change to baseline in MAP post *Pinch* in the group without lidocaine (*ED18 w/o Lido Pinch)* compared to *ED18 w/o Lido Touch, ED18 w/ Lido Touch* and *ED18 w/ Lido Pinch*. Displayed as the mean ± standard deviation. One-way ANOVA (normally distributed); p values shown. **b** Percent change to baseline in HR post *Pinch* in the group without lidocaine (*ED18 w/o Lido Pinch)* compared to *ED18 w/o Lido Touch, ED18 w/ Lido Touch* and *ED18 w/ Lido Pinch*. Displayed as the mean ± standard deviation. Kruskal− Wallis test (not normally distributed); p values shown. **c-d** Percent change from the baseline mean value in MAP and HR post *Pinch* over time; values recorded every four seconds for one minute before and one minute after stimulation (*ED18 w/o* or *w/ Lido Pinch*); values shown as the mean ± standard deviation (shaded).

The changes in HR in response to *Pinch* were slightly reduced by the application of lidocaine. However, a significant difference in HR was observed only between *ED18 w/o Lido Pinch* and *ED18 w/ Lido Touch* (p=0.0097). In the group treated with lidocaine (*ED18 w/ Lido*), no embryo showed a hyperacute decrease in HR of -15 % with a subsequent increase of 5 % from the baseline mean value post stimulus, while this reaction pattern was observed in 80 % of the embryos in the group *ED18 w/o Lido Pinch*. However, a slight decrease in HR post *Pinch* was also observed in the local anesthetic group (*ED18 w/ Lido Pinch*), but this could not be distinguished from physiological variations in HR (Fig. 3).

## Discussion

This study successfully developed methods to record blood pressure and HR in chicken embryos between EDs 7 and 18. Cardiovascular changes in response to a mechanical noxious stimulus at the base of the beak were investigated with the aim of identifying the onset of nociception during embryonic development in chicken.

While there are many well-established noninvasive methods for determining HR in the chicken embryo^24-26^, the direct intraarterial measurement is the gold standard for recording blood pressure^27^. In the past, blood pressure in chicken embryos was measured using glass capillaries or needle catheters inserted into an embryonic artery^28-30^. Corresponding to prior descriptions in the literature^28-30^, an increase in arterial blood pressure with increasing age of the embryos was observed in the present study, while there were no major differences in HR between the EDs. Thus, the optical measurement of arterial blood pressure and HR with a microtip catheter represents a reliable method for invasive measurement of blood pressure and HR in chicken embryos. However, insertion of the catheter was particularly challenging at ED7 and ED18 due to the small size of the chorioallantoic vessels at ED7 and the beginning regression of the chorioallantoic vessels at ED18.

Since self-reporting is not possible, it is difficult to evaluate pain perception in animals^22^. Nociceptive reactions to a noxious stimulus, on the other hand, can be measured^13^. The recording of cardiovascular parameters is well suited for the clinical evaluation of nociception in animals, including birds^31,32^. In the present study, the acquisition of cardiovascular parameters could be established for chicken embryos between EDs 7 and 18. Blood pressure and HR are mainly influenced by the autonomic nervous system^33^. Transmission of a noxious stimulus to the central nervous system results in activation of the sympathetic nervous system, which usually leads to an increase in blood pressure and HR^33^. Therefore, recording cardiovascular variables is considered the gold standard for the detection of nociception under anesthesia^34^.

As a means of assessing a cardiovascular response of the chicken embryo to a mechanical noxious stimulus at the base of the beak, the MAP was found to be the most sensitive parameter in the present study. Significant differences in MAP between *Pinch* and *Touch* were earliest detected on ED16, while significant changes in HR were observed only in ED17 and ED18 embryos. While there was a distinct increase in MAP in response to *Pinch* that reached over 10 % deviation from baseline in ED17 and ED18 embryos, the changes in HR were variable, and not necessarily were there any associations between changes in MAP and HR. Similar observations have been reported in adult chickens^35^. MAP has also been described in other studies concerning nociceptive responses in mammals as the most sensitive indicator of nociception^33,36^.

A prerequisite for a cardiovascular response to external stimuli is functional regulation of the cardiovascular system. Blood pressure in the chicken embryo is mainly regulated by the sympathetic nervous system^37^. An adrenergic tone on the cardiovascular system is considered to be present from a point in time halfway through the incubation period^38,39^. Therefore, the sympathetic influence on blood pressure is expected to be functional from approximately ED10^38^. In the heart, adrenergic and cholinergic receptors are already functional on ED4^40^. Changes in HR due to alterations in environmental conditions such as oxygen levels and temperature have already been observed on ED3^41^. In the present study, significant changes in HR after a noxious stimulus were not observed until ED17.

Another prerequisite for the assessment of a nociceptive response is functioning stimulus transmission. Despite some differences in the nervous system, the processing of noxious stimuli in birds is comparable to that in mammals^13^. C-fibers and A-delta fibers have been found in chickens, innervating the beak, nasal and buccal mucosa as well as the legs^11,42^. High-threshold mechanothermal nociceptors are polymodal and respond to mechanical lesions, elevated temperatures and chemical insult^13^. It is considered that injuries to the beak can be highly painful for the bird^42^ since the beak tips are intensely innervated areas^43^ and both the upper and the lower beak contain nociceptors^44^. Reflective reactions such as movements of the head to mechanical and thermal stimuli and to needle punctures appear for the first time in the skin area of the beak on ED7^45^. Therefore, in the present study, the application of a noxious stimulus to the base of the beak was chosen to evoke the highest possibility for a nociceptive response.

Regarding HR, irregularities appeared spontaneously over the whole measurement period, even at baseline. Mainly short decelerations in HR were observed, whereas MAP was not affected. It has already been reported in several publications that HR irregularities physiologically occur at the end of the second week of incubation^46-49^. Nevertheless, the HR irregularities did not have a great influence on the calculation of the mean. Minor changes in DAP reflected the HR irregularities, but the analysis of the MAP was not affected. In contrast to physiological variations in HR, a hyperacute decrease in HR with a subsequent increase could be clearly identified as a response to *Pinch* in 30 % of ED17 and 80 % of ED18 embryos. This reaction pattern could be distinguished from physiological variations in HR by the finding that post *Pinch*, HR decreased by at least -15 % and was followed by a sustained increase in HR of at least 5 % from the baseline mean value. The decrease in HR post *Pinch* was also accompanied by a short decrease in the MAP followed by an increase. A decrease in HR as a reaction to a noxious stimulus has been reported in adult chickens^35^ and in mammals^50,51^ and may be due to a vasovagal reflex to a noxious stimulus^52^. However, only single embryos showed a hyperacute decrease in HR post *Pinch*, which shows that the response in HR to a noxious stimulus is individual. Variable responses in HR after a noxious stimulus have also been described in adult chickens^35^.

In addition to a nociceptive response, it must also be considered that the measured cardiovascular changes may be induced by other factors that influence the autonomic nervous system^53^, or by embryonic movements. Especially in birds, physiological variables can be influenced by many external factors such as temperature, light or handling^53^. A correlation between fetal movements and HR irregularities has been described in human fetuses^54^. In the present study, movements of the embryo induced minor variations in HR and DAP, but the MAP was not affected. No sustained increase in MAP and HR as observed in response to *Pinch* could be attributed to movements.

Infiltration anesthesia at the base of the beak could be used to verify that the measured changes in MAP and HR may be classified as a nociceptive response and were not caused by embryonic movements or factors that influence the autonomic nervous system. The application of local anesthetics is one of the best methods to prevent the generation and transmission of nociceptive impulses^55^. The anesthetics act by blocking sodium channels in the nerve axon^53^. The application of lidocaine or bupivacaine has been described as an effective method of analgesia in birds^56^. However, the time of onset of action and the duration of action are not defined for birds^53^. In the present study, lidocaine was used because it has a rapid onset of action in mammals^55^, and in chicken, a short onset of action has also been described in spinal anesthesia^57^. Given that higher sensitivity to local anesthetics is expected in birds than in mammals^58^, embryos were intensively monitored for the occurrence of toxic effects. No signs of side effects such as bradycardia, arrythmia or hypotension were observed in the tested embryos. Since the increase in MAP was significantly reduced by the injection of lidocaine, the cardiovascular reactions to *Pinch* in the embryos that did not receive local anesthesia might be interpreted as a nociceptive response to the noxious stimulus. A limitation and possible explanation for not entirely suppressing the reaction in all embryos was that injection into the beak of the moving embryo was challenging and infiltration of the entire beak area could not always be assured. Since the present study is explorative, further investigations would need to be performed to ultimately exclude other factors as the cause of the measured cardiovascular changes. However, assuming that it is a nociceptive response, further studies regarding anesthesia and analgesia protocols are necessary to provide improved animal welfare for chicken embryos in research. Cardiovascular variations are commonly used to determine the need for analgesia or sedatives^23^ and so far there are no EU-wide regulations regarding anesthesia and analgesia for chicken embryos in research.

Although no significant difference between *Pinch* and *Touch* was reached at ED15 in MAP and HR, individual responses could be observed. Occasionally, embryos at ED15 showed reactions in MAP and HR post *Pinch*. The measurements of these embryos were performed late in the day. The development of the embryos could therefore have been more advanced compared to embryos examined in the morning. In addition, embryonic development can be influenced by various factors, and some embryos might progress faster in development than others^38^. Therefore, it has to be assumed that a nociceptive cardiovascular response is possible in individual embryos at ED15.

A limitation of the study was that intra-arterial measurement of blood pressure and HR is an invasive method. The measurements had to be performed on the opened egg, and it was inevitable to open the egg membranes. Since chicken embryos are highly sensitive to external factors^28,41^, special care was taken to maintain standardized environmental conditions and to avoid blood loss during preparation. In some embryos, severe bradycardia and hypotension were observed, or HR frequently decreased to zero. These embryos had to be excluded from the analysis because reliable measurements could not be completed. At ED7, reaching the beak was challenging, and a measurement could only be performed in 3 embryos; severe arrythmias affecting the MAP were observed. The microtip catheter is designed to measure low pressures, but the measurement accuracy of 2 mmHg, according to the manufacturer, reached its limits at the extremely low blood pressure on ED7. The results from ED7 should therefore be interpreted with caution.

## Conclusion

In conclusion, significant differences in a cardiovascular response to a mechanical noxious stimulus at the base of the beak compared with a light touch at the base of the beak were detected in chicken embryos on EDs 16 to 18. For individual embryos, a cardiovascular response to a mechanical noxious stimulus is considered to be already possible on ED15. Infiltration anesthesia with the local anesthetic lidocaine 2 % could significantly reduce reactions in MAP to a mechanical noxious stimulus at the base of the beak in ED18 embryos, indicating that the measured cardiovascular changes can be interpreted as nociceptive responses.

## Material and Methods

### Animals

Fertilized eggs from a breeding of Lohman Selected Leghorn chicken eggs were obtained from the TUM Animal Research Center (Thalhausen) and stored at 15 °C. ED0 was considered the day when eggs were transferred to the incubator (Favorit – Olymp 192 Spezial, HEKA – Brutgeräte, Rietberg, Germany). The eggs were incubated for 7 to 18 days at 37.8 °C and 55 % humidity and turned six times a day until they were fenestrated.

At ED3 of incubation, the egg shells were fenestrated^59^. For this purpose, the egg was placed horizontally for at least two minutes, and then 5 to 7 ml albumen was withdrawn from the apical pole of the egg using a 5 ml syringe and an 18 G needle. The top of the egg was then covered with tape. A hole was cut in the shell, and the vitality of the embryo was verified. Then, 0.5 ml penicillin-streptomycin (10,000 units penicillin, 10 mg streptomycin/ml, P4333-100 ml Sigma−Aldrich) was added before the egg was resealed with cling film and was further incubated in horizontal position. The vitality of the embryos was checked daily until the end of the experiment. Experiments were conducted between 9:00 a.m. and 7:00 p.m. so that the variance in the age of the embryos within an ED was limited to a maximum of 10 hours.

## Experimental design

The study was an explorative study. For ED12 to ED18, n=10 embryos of each embryonic day were measured. Due to higher losses in younger embryos, a group size of n=6 (ED9) and n=3 (ED7) embryos was chosen. Furthermore, to study the effect of local anesthesia, n=6 ED18 embryos were used.

Experiments were performed under standardized conditions in a specially designed heating chamber equipped with a heating lamp (ARTAS GmbH, Arnstadt, Germany) and an air humidifier (HU4811/10 Series 2000, Philips, Amsterdam, the Netherlands). The eggs were placed on a heating mat (ThermoLux, Witte + Sutor GmbH, Murrhardt, Germany) in a bowl filled with warmed Armor Beads (Lab Armor Beads™, Sheldon Manufacturing, Cornelius, USA). The mean temperature and mean humidity during all experiments were 37.7 °C ± 0.8 and 55.5 % ± 4.3.

First, the cling film was removed from the egg, and the shell was carefully opened to the level of the chorioallantoic membrane. Using a microscope (Stemi SV6, Zeiss, Oberkochen, Germany), the allantoic and amniotic membranes were opened over the head of the embryo, avoiding any large vessels so that the beak could be reached in the further course of the experiment. A side branch of the chorioallantoic artery was prepared, temporarily ligated to avoid blood loss, and incised with microsurgical scissors. A microtip catheter (FISO-LS Fiber Optic Pressure Sensor, FOP-LS-PT9-10, FISO Technologies Inc., Quebec, Canada) was then inserted into the vessel and fixed in place with a ligature. SAP, DAP, MAP and HR were recorded continuously every four seconds (PLUGSYS module, EIM-B, EIM-A, HAEMODYN Software, Hugo Sachs Elektronik-Harvard Apparatus GmbH, March-Hugstetten, Germany, FFP-LS and Evolution Software, FISO Technologies Inc., Quebec, QC, Canada). The beak of the embryo was carefully placed on a Desmarres lid retractor. For younger embryos at ED7 and ED9, the beak was carefully placed on a custom-made wire loop.

After implementation of the catheter, a two-minute waiting period followed. Then, two mechanical stimuli were applied at the base of the beak. In randomized order, either a mechanical noxious stimulus was applied with a surgical clamp (*Pinch*) or, as a negative control, a light touch (*Touch*) was applied first. Between the two stimuli, a period of five minutes was maintained to normalize the parameters. After the second stimulus, measurements were continued for five more minutes. The measurement time between both stimuli and after the second stimulus was reduced from five to three minutes for ED13 and younger embryos due to the increasing sensitivity of the organism.

For the *Pinch*, a surgical clamp was applied to the base of the beak and squeezed. For the *Touch*, the beak was only lightly touched with the surgical clamp. For both stimuli, a mosquito clamp was used for ED12 to ED18 embryos. For embryos at ED7 and ED9, the surgical clamp was too large, and microsurgical forceps were used instead for both stimuli. To ensure comparability, the stimuli were always applied by the same person. In the further course of the study, an analgesia meter (BIO-RP-M, BioSeb, Vitrolles, France) with customized tips of the mosquito clamp was used to monitor the pressure applied by the mechanical stimulus.

To verify whether the measured cardiovascular responses could be classified as nociceptive responses, in n=6 ED18 embryos, a local anesthetic was applied before stimulation. For this purpose, after the preparation and placement of the microtip catheter, 0.02 ml of lidocaine 2 % (Xylocitin® 2 %, Mibe GmbH Arzneimittel, Brehna, Germany) was injected into the upper and lower beak using a 30 G needle (*ED18 w/ Lido Touch* and *Pinch*). The measurements were carried out following the same experimental protocol as for ED14 to ED18 embryos with the exception that a waiting period of three minutes was added prior to the measurement. During this time, blood pressure and heart rate were monitored for the occurrence of side effects of lidocaine, such as bradycardia, arrythmia or hypotension. As a comparison group without lidocaine, the already measured ED18 embryos were used (*ED18 w/o Lido Touch* and *Pinch*).

Immediately after the end of the experiments, the embryos were euthanized by intravenous injection of pentobarbital sodium (Narcoren®, 16 g/100 ml, ED7-ED12: 0.1 ml; ED13-ED19: 0.2 ml) followed by decapitation.

### Analysis

SAP, DAP, MAP and HR were recorded every four seconds. For the evaluation of the reactions to the stimuli, the means of MAP and HR were calculated over one minute before (= baseline) and one minute after the respective stimulus. To avoid an influence by the approach of the clamp, the 15 seconds immediately before the respective stimulus were not included as part of the baseline. In embryos showing a hyperacute decrease in HR with a subsequent increase in HR post *Pinch*, the decrease was not included in the calculation and was evaluated separately to avoid compensation of opposite reactions. The percentage deviation of the response after the stimulus (*Pinch/Touch*) to the baseline was then calculated. Differences in the percent changes to baseline in MAP and HR after *Pinch* and *Touch* were tested for statistical significance. For normally distributed data, a paired t test (two-tailed) was used. For data that failed the normality test, a Wilcoxon signed-rank test (two-tailed) was performed. For the comparison of multiple groups, either a one-way ANOVA (normally distributed) or a Kruskal−Wallis test (not normally distributed) was used.

## Data availability

Raw data are available up on reasonable request to the corresponding author.

## Acknowledgments

This work was financially funded by the German Federal Ministry of Food and Agriculture (BMEL) based on a decision of the Parliament of the Federal Republic of Germany, granted by the Federal Office for Agriculture and Food (BLE; grant number 2821HS005).

The authors thank the scientific advisory board with Prof. Dr. Dr. Michael Erhard, Prof. Dr. Wolf Erhardt, Prof. Dr. Harald Luksch, Prof. Dr. Heidrun Potschka, Prof. Dr. Hans Straka and Dr. Britta Wirrer for their excellent scientific contribution as well as Dr. Johannes Fischer, Stefanie Fitzner and Dr. Hicham Sid for their technical support.

## Author contributions

Conceptualization: CB, AMS, JW, JR, TF, BS; Data curation: LW, AMS; Formal analysis: LW, AMS, JW; Funding acquisition: CB, TF, BS; Investigation: LW, SCS, AMS, JW, JR; Methodology: CB, AMS, JW, JR, SK, MA, TF, BS; Project administration: CB; Resources: CB, BS; Supervision: CB, TF, BS; Writing - original draft: LW; Writing - review & editing: LW, AMS, JW, SCS, JR, SK, MA, TF, BS, CB. All authors have read and agreed to the published version of the manuscript.

