## Supplementary Information for "Nociception in chicken embryos, Part I: Analysis of cardiovascular responses to a mechanical noxious stimulus"

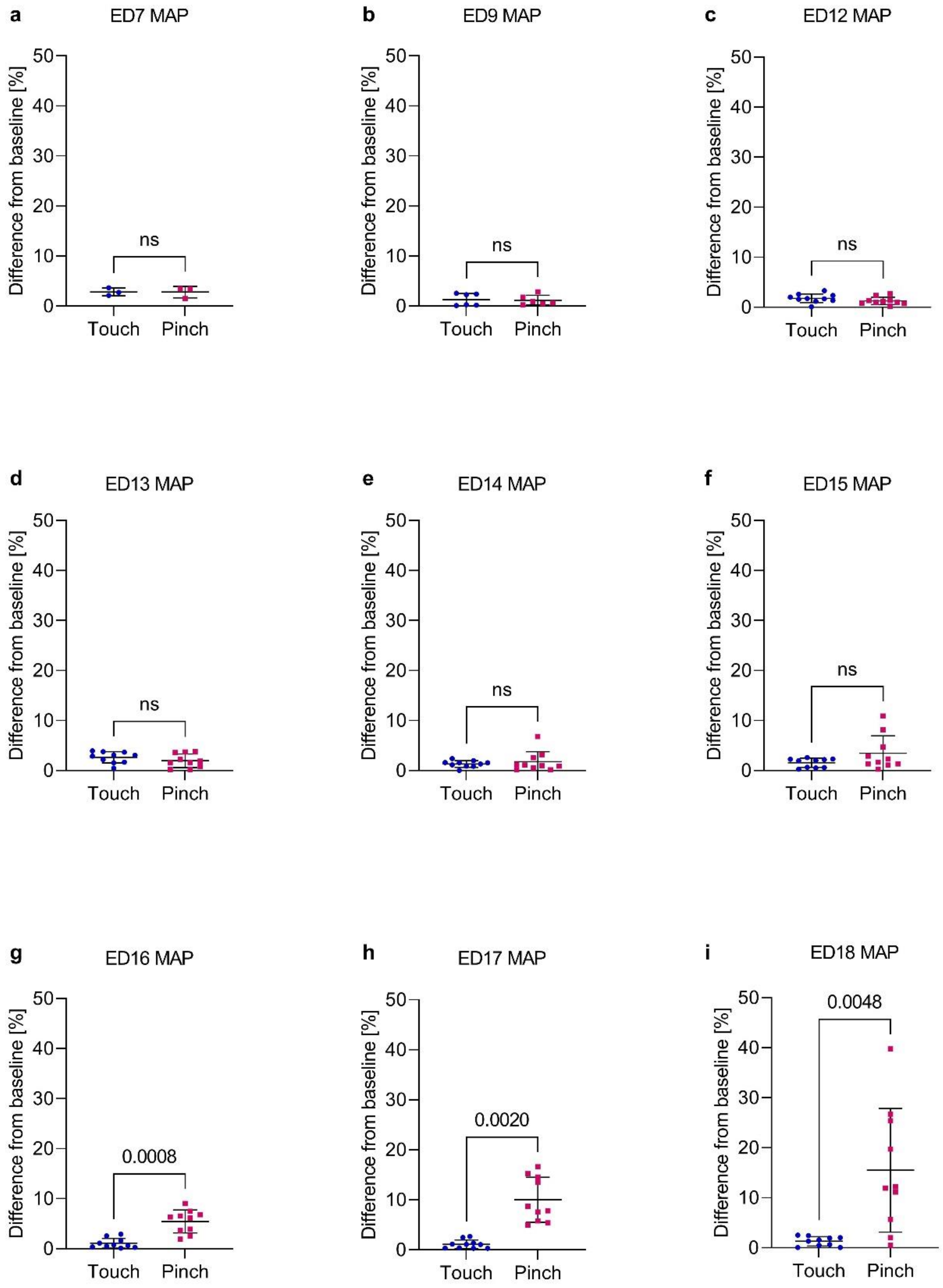


**Supplementary Fig. 1. Percent change in MAP post *Touch* and *Pinch*. a-i** Embryos at embryonic day (ED) 7 (n=3), ED9 (n=6) and EDs 12 to 18 (n=10) received a mechanical noxious stimulus (*Pinch*) and a light touch as control (*Touch*) at the base of the beak in randomized order. Displayed as the mean ± standard deviation. Paired t test (normally distributed: c, d, g and i) or Wilcoxon signed-rank test (not normally distributed: a, b, e, f and h). P values shown; g: p=0.0008, h: p=0.0020 and i: p=0.0048; ns = no significant difference between the groups.

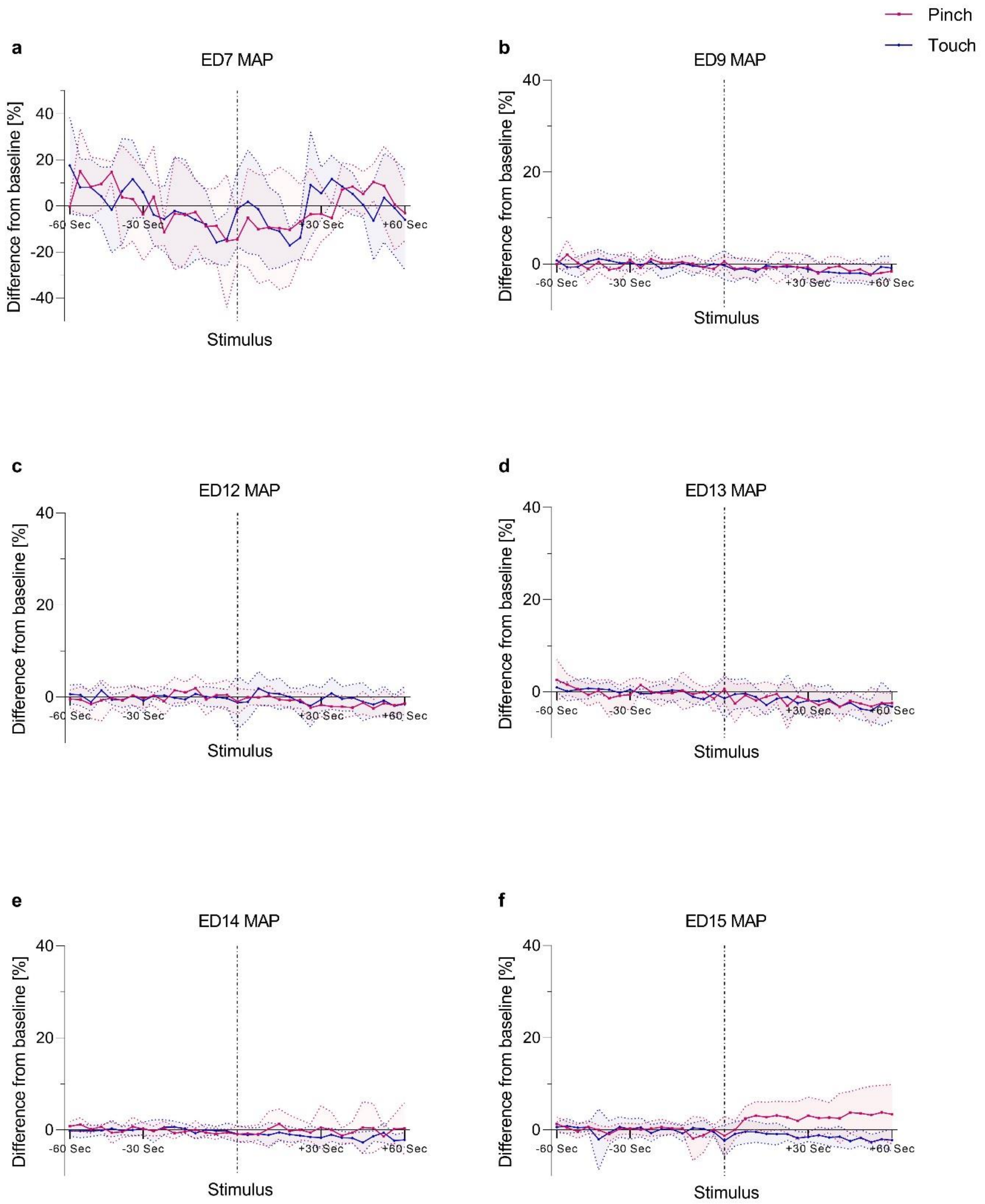


**Supplementary Fig. 2. Percent change from the baseline mean value in MAP over time. a-f** Embryos at ED7 (n=3), ED9 (n=6) and EDs 12 to 15 (n=10) received a mechanical noxious stimulus (*Pinch*) and a light touch as control (*Touch*) at the base of the beak in randomized order. Values were recorded every four seconds for one minute before and after stimulation (*Touch* and *Pinch*). Values are shown as the mean ± standard deviation (shaded).

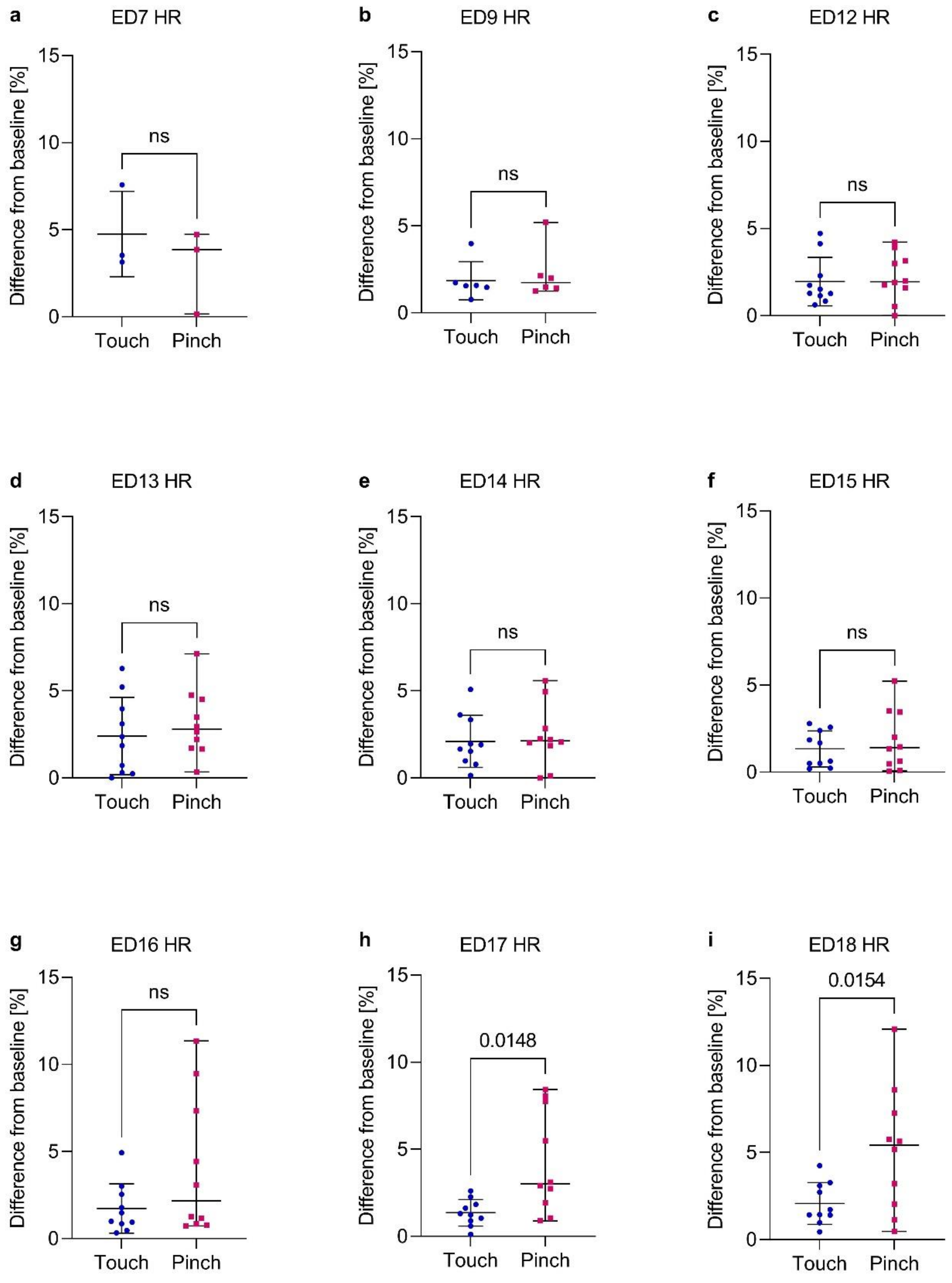


**Supplementary Fig. 3. Percent change in HR post *Touch* and *Pinch*. a-i** Embryos at ED7 (n=3), ED9 (n=6) and EDs 12 to 18 (n=10) received a mechanical noxious stimulus (*Pinch*) and a light touch as control (*Touch*) at the base of the beak in randomized order. Displayed as the mean ± standard deviation. Paired t test (normally distributed: a, d-f, h and i) or Wilcoxon signed-rank test (not normally distributed: b, c and g). P values shown; h: p=0.0148 and i: p=0.0154; ns = no significant difference between the groups.

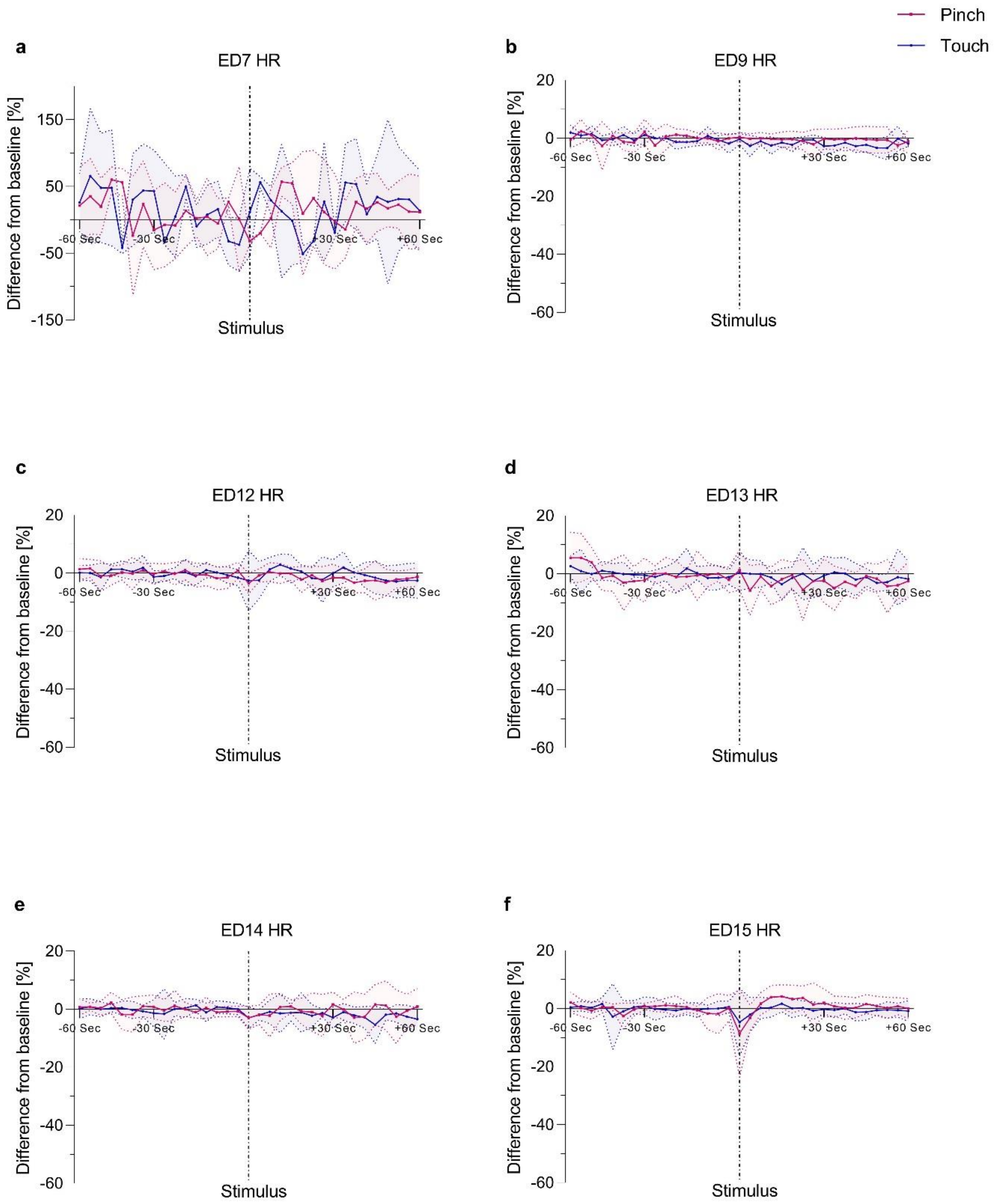


**Supplementary Fig. 4. Percent change from the baseline mean value in HR over time. a-f** Embryos at ED7 (n=3), ED9 (n=6) and EDs 12 to 15 (n=10) received a mechanical noxious stimulus (*Pinch*) and a light touch as control (*Touch*) at the base of the beak in randomized order. Values were recorded every four seconds for one minute before and after stimulation (*Touch* and *Pinch*). Values are shown as the mean ± standard deviation (shaded).
